# The Stop Signal Stepping Task: how action cancellation commands disrupt step initiation in young and healthy older adults

**DOI:** 10.1101/2025.02.24.640005

**Authors:** Rebecca Healey, Marlee J. Wells, Sauro E. Salomoni, Rohan Puri, Mark R. Hinder, Rebecca J. St George

## Abstract

*Action cancellation* – the ability to rapidly cancel an initiated movement in response to unexpected events – has been extensively studied in the upper limb using the stop signal task (SST). During gait, action cancellation is needed to stop and modify steps to avoid hazards and prevent falls. By adapting the SST to step initiation, this study investigated how the anticipatory postural adjustment (APA) and foot-lift phases of forward stepping were affected by action cancellation commands, and whether this changed with healthy ageing. The SST was performed in stepping, foot tap, and finger button conditions in 27 young (*M*_age_ = 28.7 years) and 29 healthy older adults (*M*_age_ = 70.1 years).

Across conditions, older adults exhibited slower *response* speed compared to young adults and greater proactive slowing of responses when stop cues were anticipated. However, there was no significant difference in *stopping* speed between young and older adults. Stopping speed was fastest in the finger tap condition, and slowest in the step condition. When an APA was initiated in a step cancellation trial, the magnitude of the weight shift toward the step leg did not differ between successful and unsuccessful foot-lift cancellations. Foot-lift could be cancelled when stop cues were presented at similar phases of step preparation for young and older adults.

These results suggest that the initial loading of the step leg is a ballistic process, however as weight is shifted toward the stance leg, action cancellation commands responding to external stimuli can decouple the APA and foot-lift step phases.

**Key Points:**

- The stop signal task (SST) – which allows an estimation of stopping speed independently of response speed – was applied to voluntary stepping in young and older adults.
- While response speed was slower for older than young adults, stopping speed was not significantly different between age groups in the upper limb, lower limb when seated, and during forward stepping.
- When stop cues were introduced, response speed slowed more in older than young adults, and more in the upper than the lower limb (i.e., Foot Tap and Step conditions).
- The initial preparatory weight shift toward the stepping foot was not significantly different between successfully cancelled steps and normal steps, highlighting the ballistic nature of the early phase of step preparation.
- Prior to foot-lift, action cancellation commands could decouple the preparatory weight shift phase from foot-lift at similar stages of step initiation in young and healthy older adults.

## Introduction

Imagine standing at a busy intersection, waiting to cross the road. The green ‘walk’ symbol flashes, prompting you to step forward. Suddenly, you notice an oncoming car has ignored the red traffic light, forcing you to quickly stop your step to avoid a collision. In daily life, the ability to rapidly cancel or modify initiated actions in response to unexpected sensory information is a vital executive function. Despite the importance of inhibitory control for adaptive stepping movements, the gold standard laboratory test of response inhibition – the stop signal task (SST) – has almost exclusively been applied to upper limb responses while seated. In a typical SST, participants perform a choice reaction time (CRT) task, responding as quickly as possible to ‘go’ signals (e.g., left- or right-pointing arrows) by pressing buttons. On a small percentage of trials, a ‘stop’ signal appears shortly after the ‘go’ signal, indicating participants should try to cancel their initiated response. By manipulating the temporal delay between the ‘go’ and ‘stop’ signals so that successful stopping approximates 50%, the speed of stopping (the stop signal reaction time – or SSRT) can be estimated independently of response execution speed ^1^.

A growing body of evidence suggests that response *execution* speed and response *stopping* speed may be independent processes that are differentially affected by ageing. In simple and choice reaction time tasks – where quick responses are crucial, and the possibility of having to stop is *not* part of the response set – healthy older adults consistently exhibit slower response *execution* speed compared to young adults for both upper limb ^2^ and stepping tasks ^3^. However, SST studies in the upper limb indicate that the impact of ageing on *stopping* speed is less consistent. While some studies report age-related slowing of SSRTs ^4–6^, others find no significant differences between young and older adults ^7^, or report that the rate of age-related slowing in stopping speed is considerably less than the rate of slowing in response execution speed ^8,9^.

Response execution speed to ‘go’ trials within stop signal paradigms (representing the majority of trials) tend to be slower than during equivalent CRT tasks (when stopping is not anticipated) and this response slowing is greater in older, compared to young adults ^10^. Thus, ‘go’ responses during SSTs reflect not only motor execution capacity (or speed) but also strategic processes related to the uncertainty of the situation, as stopping might be needed. One such strategy is *proactive slowing,* which refers to a slower, more cautious approach to initiating a response ^11^, conceivably to increase the chances of successful cancellation *if* a stop signal does occur ^12^. The degree of proactive slowing can be determined by comparing response times during ‘go’ trials in SSTs to those in a commensurate CRT task that lacks the possibility of stopping ^11^. However, a more detailed approach involves examining response slowing in the subsequent ‘go’ trials immediately following stop trials ^13,14^ or specifically after unsuccessful stopping trials ^5,15–17^. It is unknown whether proactive slowing occurs in stepping tasks under uncertain conditions that may require sudden step cancellation.

Inhibitory control ability has been linked to falls risk in older adults^18,19^. This is thought to be due to the role of inhibitory control in gait adaptation – that is, the ability to modify a planned step in line with changing environmental demands. People with fast SSRT on a seated paradigm make more adaptive changes to the velocity and joint angle of the ankles and knees during an unplanned gait termination task^20^ and are more able to inhibit unwanted leg movements in a reactive balance paradigm, in favour of a handle grasping response^21^. However, these studies used correlational designs and did not measure ability to inhibit an involuntary step in the absence of balance perturbation.

Previous research has explored relationships between inhibitory control in the hands and feet using seated SSTs, suggesting longer SSRTs in the feet than the hands^22–24^. Given that lower-limb inhibitory control appears to take longer than upper-limb inhibitory control, and that inhibitory control ability appears integral to balance in older adulthood, it is of interest to explore differences in how young and older adults use inhibition during standing balance. Recently, there has been a push to translate seated upper-limb inhibitory paradigms to standing balance paradigms, in order to measure gait-based inhibition more directly^25^.

Previous studies of inhibitory control during step initiation have assessed the ability to generate a forward step in some trials but withhold a step on other trials – as in the Go/No-Go task^26^. Such paradigms measure *action restraint –* the ability to withhold an anticipated action – rather than *action cancellation*, which involves cancelling an already initiated action. This is an important distinction, as recent neurophysiological work suggests that these processes operate via different neural mechanisms^27^. Older adults’ inhibitory stepping ability has also been assessed using adaptions of the Stroop Task^26,28,29^, Simon Task^29–33^, and Flanker Task^33–35^. However, it can be argued that these paradigms assess *perceptual inhibition* and *cognitive inhibition* – that is, the ability to ignore irrelevant or confusing visual stimuli during step decision making, and the ability to override a dominant response, respectively. Whilst they are very informative when investigating step preparation and complex decision making, they are less able to inform on how the ability to cancel an initiated step changes with age.

A recent study demonstrated feasibility of exploring step cancellation ability in young and older adults during an adapted stop signal task^36^. However, due to low stop trial numbers (total 9 per participant), high rate of data exclusion (7% of trials in young adults, and 16% in older adults), and low rate of successful stopping (29% in young, and 19% in older adults), a detailed understand of the interaction between inhibitory control and postural control was not possible. In a similar vein, researchers have assessed response inhibition during steady gait by having participants step onto targets projected on the ground, while attempting to avoid the targets if they changed colour mid-step ^37^. This study reported that older adults failed to modify steps more often than young adults. However, as this study used a fixed time interval between the presentation of the target stimulus and the avoidance cue, it could not isolate stepping stopping speed from response speed. Thus, it remains unclear whether older adults have a specific deficit in the time required to cancel initiated steps, and at what stage a step *can* be cancelled in young and older adults.

Stepping is inherently more complex than seated finger button presses. Before the stepping foot leaves the ground, characteristic changes in ground reaction forces (ΔGRFs) – detectable with force plates under the feet – occur, shifting the centre of pressure by loading and then unloading the stepping leg to facilitate subsequent movement of the centre of body mass forward and toward the stance leg ^38^. Known as Anticipatory Postural Adjustments (APAs), this behaviour is thought to operate under a feedforward, ballistic mode of control ^39,40^ and contribute to postural stability during voluntary movements. The APA (pre-foot-lift weight shifts) and swing (foot-lift to heel strike) phases of the step are intrinsically linked, with the speed and direction of the APA determining the location of the final stepping target^41^. Despite this close coupling, the APA and foot lift phase can become uncoupled in some circumstances. For example, people with freezing of gait in Parkinson’s Disease are generally able to execute an APA but are unable to trigger the foot lift ^42,43^. The current study used the SST to investigate the extent to which volitional action cancellation commands responding to visual cues can interrupt the coupling between the APA and foot-lift phases of the step. If APAs are a ballistic process, resistant to volitional commands once set in motion, the profile of ground reaction forces on ‘successful’ stop trials (i.e., an APA was initiated in response to the ‘go’ cue, but the foot lift was cancelled following the ‘Stop’ cue) will be comparable to the profile on stepping trials.

The study also sought to determine whether healthy ageing affects the speed at which a volitional step can be cancelled. It was hypothesised that due to the greater complexity of stepping actions and the need to control balance, stopping a step would take longer than stopping either a seated foot or finger response, and that due to age-related balance impairment^44^, stopping a step would be slower in older compared to young adults.

## Materials and Methods

### Participants and Ethical considerations

This study was approved by the University of Tasmania Human Research Ethics Committee (approval code: H0014865) and preregistered on the Open Science Framework website (https://osf.io/ymvsu/). Signed, informed consent was obtained from all participants in accordance with the Declaration of Helsinki.

A convenience sample of 56 participants, with young (n= 27; age range: 18-43 years) and older (n= 29; age range: 60-79 years) adults recruited from the University of Tasmania and broader local community, participated in the study (demographic characteristics are presented in Table 1). Exclusion criteria included neurological conditions (e.g., history of stroke, Parkinson’s Disease, or dementia), past neurosurgery, and pain during standing and walking. Older adults had no cognitive impairment (standardised Mini Mental State Exam [sMMSE] score >27). Data were excluded for two participants in the young group on the Foot Tap and Step conditions due to technical issues or early experiment termination. Participants reported the number of falls in the past year, and older adults additionally completed the Falls Self-Efficacy Scale (FES-1; see Table 1) ^45^. Participants who reported one or more falls over the past 12 months were classified as “fallers”. A Chi square test of independence indicated that the number of fallers did not differ by Age Group, χ^2^ (56) = 1.03, *p =* .310.

**Table 1.**
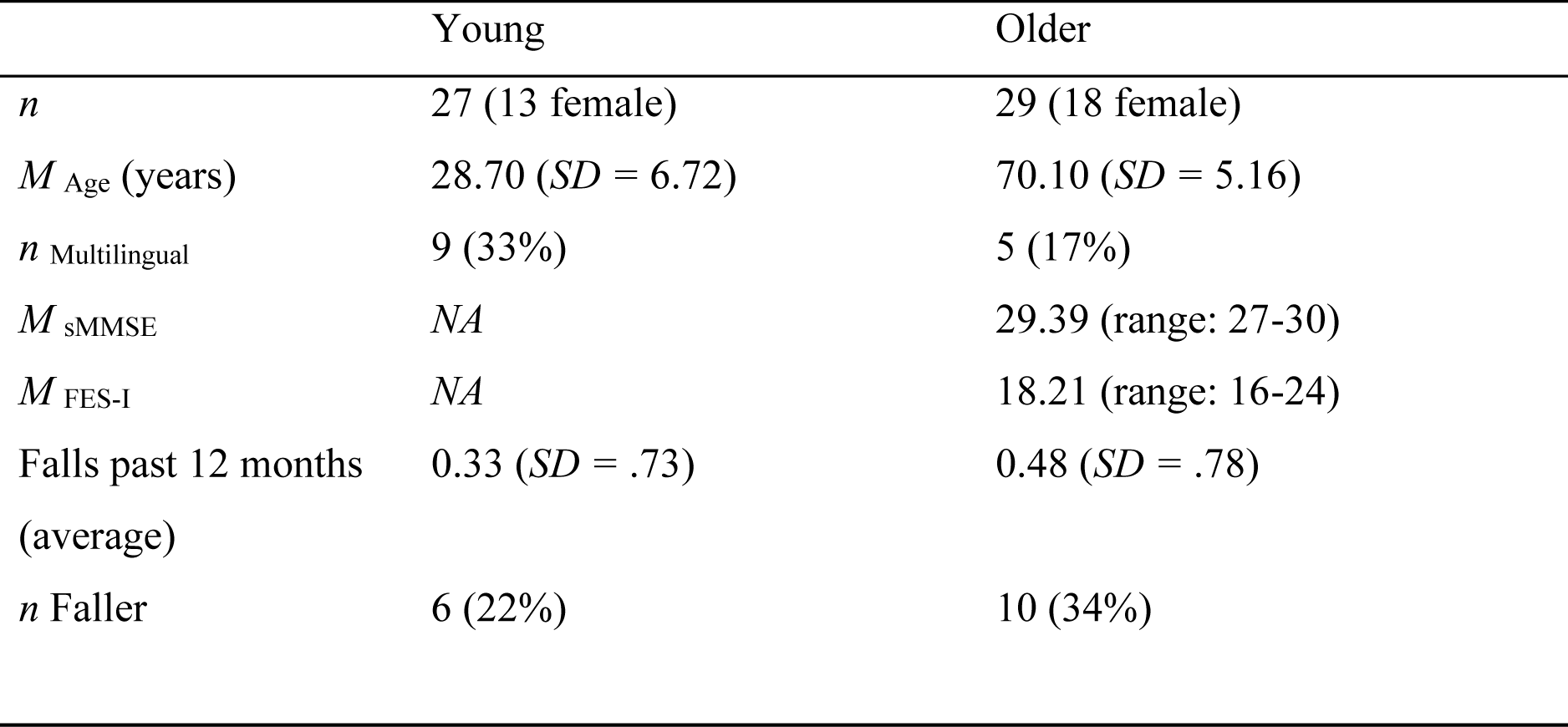
Characteristics of the Participant Samples.

Characteristics of the Participant Samples
|  | Young | Older |
| --- | --- | --- |
| *n* | 27 (13 female) | 29 (18 female) |
| *M*<sub>Age</sub> (years) | 28.70 (*SD* = 6.72) | 70.10 (*SD* = 5.16) |
| *n* Multilingual | 9 (33%) | 5 (17%) |
| *M*<sub>sMMSE</sub> | NA | 29.39 (range: 27-30) |
| *M*<sub>FES-I</sub> | NA | 18.21 (range: 16-24) |
| Falls past 12 months<br>(average) | 0.33 (*SD* = .73) | 0.48 (*SD* = .78) |
| *n* Faller | 6 (22%) | 10 (34%) |

Multilingual participants completed the Language Experience and Proficiency Questionnaire ^46^. Multilingualism is thought to influence inhibitory control in older adulthood ^47,48^. As such, participants self-reported the number of languages they could speak to proficiency (*n* _Monolingual_ = 42; *n* _Multilingual_ = 14). A Chi square test of independence indicated that categorisation as “monolingual” and “multilingual” did not depend on Age Group, χ^2^ (56) = 1.93, *p* = .165.

### Experimental Design

Participants completed three conditions in a counterbalanced order, all of which used a stop signal reaction time task to measure response execution and stopping speed, but which varied in regard to the effector used, and whether the participant was standing or sitting: (1) Finger Tap, where they were sat at a computer desk and the speed of index finger button presses was measured (Fig1A); (2) Foot Tap, a seated condition that assessed the speed of raising the foot off the ground (Fig1B), (3) Step initiation from quiet standing, which measured the speed of the first foot lift off the ground when transitioning the body forward i.e., there were two steps required to complete the task so that APAs were elicited in ground reaction forces (Fig1C). Task design is depicted in Table 2. All participants were provided with regular breaks within and between conditions to prevent fatigue. Young adults generally completed data collection in a single 2.5-hour block. Older adults, who required more time due to completion of the additional questionnaires, generally completed data collection over two 1.5-hour sessions, with at least 24 hours between sessions to minimise fatigue.

**Table 2.** Task Design.

Task Design
| Session (by Age) | | Block Type | | Condition ($n_{\text{blocks}}$ X $n_{\text{trials per block}}$) | | |
| --- | --- | --- | --- | --- | --- | --- |
| Young | Older |  |  | Finger Tap | Foot Tap | Step |
| Session 1 | Session 1 | CRT | $n_{\text{practice}}$ | 1 x 8 | 1 x 8 | 1 x 8 |
| Session 1 | Session 1 | CRT | $n_{\text{experiment}}$ | 1 x 32 | 1 x 32 | 1 x 32 |
| Session 1 | Session 1 | SST | $n_{\text{practice}}$ | 1 x 8 | 1 x 8 | 1 x 8 |
| Session 1 | Session 1 | SST | $n_{\text{experiment}}$ | 1 x 80 | 1 x 80 | 1 x 80 |
| Session 1 | Session 2 | SST | $n_{\text{experiment}}$ | 2 x 80 | 2 x 80 | 2 x 80 |

CRT was measured in response to ‘Go’ signals only (i.e., left- or right-facing white arrows with no stopping cues). Participants completed a block of 32 CRT trials per condition (Finger Tap; Foot Tap; and Step). Participants then completed three blocks of the SST with 80 trials each (i.e., 240 trials per condition containing both ‘Go’ and ‘Stop’ trials). Eight practice trials for each task (CRT/SST) and condition were performed but excluded from analysis. ‘Stop’ signals, indicated by the white ‘Go’ arrow turning blue, occurred randomly in 25% of SST trials, required that participants attempt to cancel their response. The delay between the ‘Go’ and ‘Stop’ cues, known as the stop signal delay (SSD), was adjusted based on stop trial success using a staircase tracking procedure. The SSD was reduced by 50ms after failed stops (to *increase* the chance of stopping successfully on the next stop trial) and was increased by 50ms after successful stops (to *reduce* the chance of stopping successfully on the next stop trial). These adjustments ensured that the overall probability of successful stopping approached 50% for each condition. SSDs were tracked independently for each condition.

A fixation cross, which prompted participants to direct their attention towards the centre of the screen, was displayed for between 0.5-1.0 second, selected randomly from a truncated exponential distribution to minimise anticipation of the ‘Go’ signal. The ‘Go’ signal appeared immediately after the fixation cross disappeared. Participants were instructed to respond as quickly as possible to the ‘Go’ signal and not to wait for the possibility of stop cues occurring. In the step condition, a walking frame was placed two steps in front of participants, to provide stability if needed, and individuals who were identified as having a higher risk of falls wore a harness suspended from the ceiling (1 older participant).

A freely available version of the SST (STOP-IT2) running in Matlab Psychtoolbox ^49^ was adapted for use in this study. The finger tap condition involved using the left and right index fingers to press the ‘F’ and ‘J’ keys of a QWERTY keyboard to respond to left and right arrows respectively. During Foot Tap and Step conditions, bilateral force plates (Accusway, AMTI, Watertown, MA, USA) recorded the ground reaction forces (GRF) and moments under each foot, sampling at 1000 Hz and low pass filtered (4^th^ order Butterworth) with a cut-off frequency of 20 Hz.

Baseline GRFs were established by initially measuring GRFs under each foot over a 1 second period for the Step and Foot Tap conditions. In the Step condition, participants stood in quiet stance, feet positioned at a comfortable width, with each foot on a force-plate. In the Foot Tap condition, they sat in a comfortable position with arms supported on armrests, knees bent at approximately 90 degrees and each foot resting flat on a force plate. In Step and Foot Tap conditions, a trial only began when the real-time weight distribution matched this baseline level to ensure posture was in a standardised neutral position at the start of each trial. For Foot Tap, the task was to tap the heel of the cued foot forwards (Figure 1B) as quickly as possible. For the Step condition, participants were instructed to step forward leading with the cued foot and following through with a second step from the trailing (un-cued) foot, to ensure the body was wholly displaced forward. Step and Foot Tap response times were identified as the point of ‘foot-lift’– when the vertical GRF of the stepping foot was less than 2% of the level at trial onset (see Figure 1).

**Figure 1.**
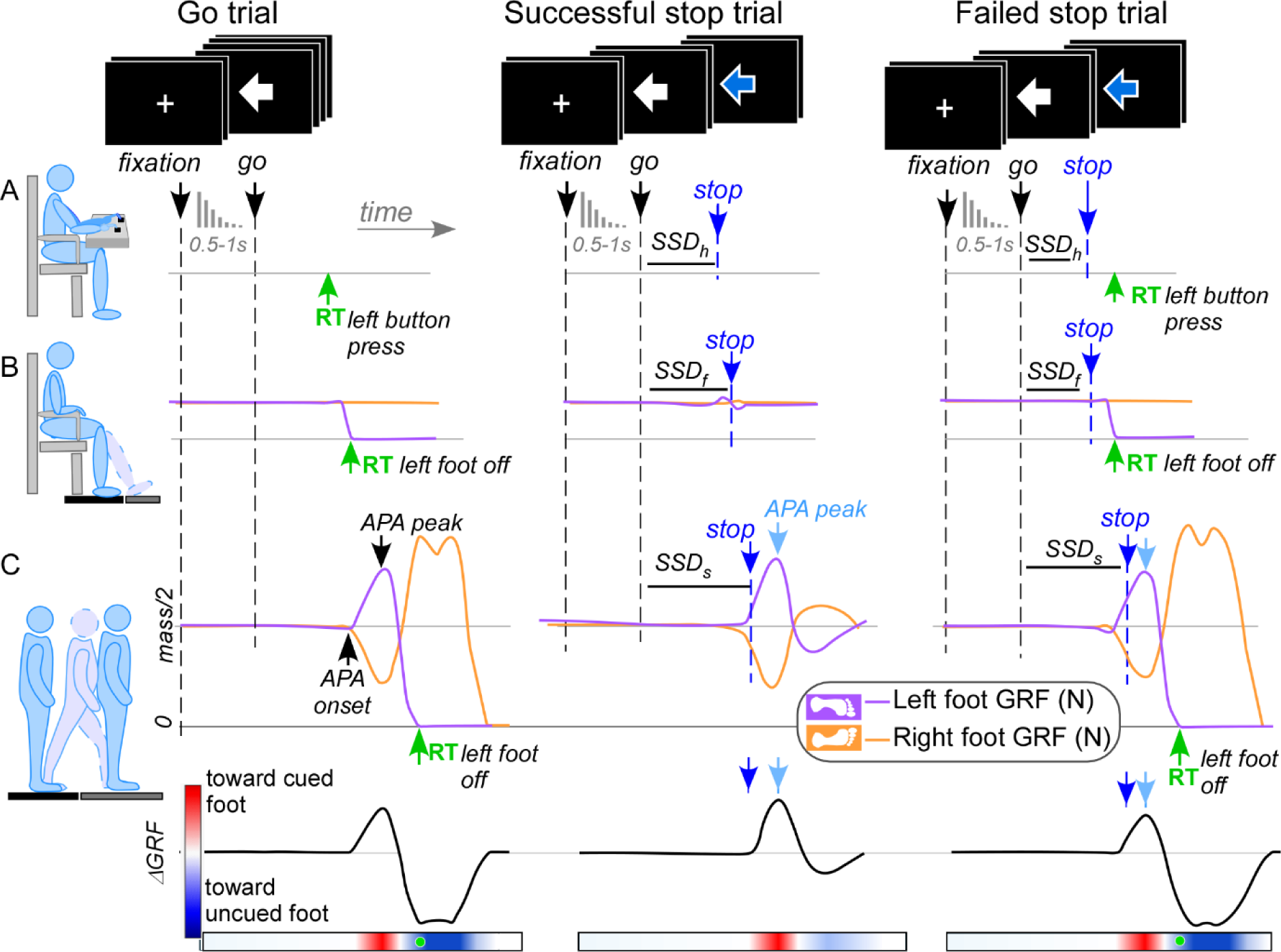
Finger Tap (A), seated Foot Tap (B), and Step (C) conditions. Typical example responses to Go only trials are shown on the graphs on the left, successful stop trials in the middle column, and failed stop trials on the right. The top of the figure (with the overlaid screen panels) shows the visual stimuli presented to participants during go and stop trials: a central fixation cue (white cross), succeeded by a go cue (white central arrow presented on both go and stop trials), and a stop cue (arrow colour change from white to blue). Pressing the button (row A) or lifting the cued foot (rows B & C) on a stop trial resulted in a failed stop. The Stop Signal Delay (*SSD*) was recalculated after each stop trial depending on the trial outcome for the Finger Tap (*SSD_h_*), Foot Tap (*SSD_f_*) and Step (*SSD_s_*) conditions. Note that for the Step condition (C), each force plate bore approximately half the total body mass (*mass/2*) before the ‘Go’ signal, and a step was initiated by firstly shifting the body weight *toward* the cued stepping leg – indicated by an *increase* in the vertical force under the cued foot and a *decrease* in the un-cued foot. The bottom row depicts the difference in ground vertical reaction force (ΔGRF waveforms) between the leading and trailing limbs in the Step condition, with the corresponding heat map of the ΔGRF magnitude shown for this example trial, with red indicating a weight shift initially toward the limb of foot-lift (in the cued direction for a correct response), blue showing a weight shift toward the trailing leg and the green dot showing the RT. The normalised heatmaps of ΔGRF for every participant and every stepping trial are present in Figure 3.

### Statistical Analysis

Data were analysed using custom MATLAB code (Mathworks Inc., Natick, MA, USA, v. r2020b) and R code (R Core Team, 2022) using the packages tidyverse (Wickham *et al.*, 2019), lme4 (Bates *et al.*, 2014), emmeans (Lenth & Lenth, 2018), matrix (Bates *et al.*, 2017), lmertest (Kuznetsova *et al.*, 2017), and multcomp (Hothorn *et al.*, 2016). All behavioural data plots were created using ggplot2 (Wickham *et al.*, 2016) and afex (Singmann *et al.*, 2015).

### Response execution speed

Go trials in the CRT and SST Blocks were filtered to remove incorrect responses, anticipatory responses (‘Go’ trial RT <150ms in the Finger Tap condition, <300ms in the Foot Tap condition, <400ms in the Step condition), and trials with no response. This resulted in the removal of 0.31% of trials, suggesting strong compliance with task instructions. The filtered dataset comprised 34,813 trials across participants, (Finger Tap = 11,784 trials [33.8%], Foot Tap = 11,380 trials [32.7%], Step = 11,649 trials [33.5%]). To quantify proactive RT slowing across the different response modalities, three different analyses were conducted, each using a generalised linear mixed model with a gamma distribution and log link function to account for positively skewed data synonymous with RT ^50^. To reduce Type I error, for each model we specify the maximal possible random effect structure that still allowed convergence ^51,52^. Significant main and interaction effects were explored using Bonferroni-adjusted pairwise tests, and marginal means are reported.

The general proactive slowing between RT of trials in the CRT block (no anticipation of stopping) to ‘Go’ trials in the SST block (some stopping anticipated) was determined with fixed factors of Block Type (CRT block, SST block), Age Group (Young, Older) and Condition (Finger Tap, Foot Tap, Step). Participant-level random intercepts were included, and slopes were allowed to vary randomly for each participant by Condition, and Block Type, and their interaction.

Response slowing for consecutive ‘Go’ trials following ‘Stop’ trials was assessed by creating a factor called ‘Trial Post Stop Signal’ (TPSS) with seven levels. Trials were categorised relative to the *most recent preceding* stop trial. Go trials occurring 1 - 6 trials after a stop signal were labelled “Post 1”, “Post 2”, “Post 3”, “Post 4”, “Post 5” and “Post 6”, respectively, with any additional trials coded as “Post 7+”. The resultant dataset comprised 29,449 trials, categorised as Post 1 trials (*n* = 7374 [25%]), Post 2 trials (*n* = 5535 [18.79%]), Post 3 trials (*n* = 4144 [14.07%]), Post 4 trials (*n* = 3050 [10.36%]), Post 5 trials (*n* = 2220 [7.54%]), Post 6 trials (*n* = 1590 [5.40%]), and Post 7+ trials (*n* = 5536 [18.79%]). This RT data was analysed using a three-way GLMM with fixed factors of Age Group, Condition, and TPSS. Participant-level random intercepts were included as the model failed to converge with random slopes.

Finally, we explored whether failed stop trials compared to successful stop trials impacted RT on the first subsequent ‘Go’ trial. A categorical variable labelled ‘Stop Outcome’ was entered into this model, with Go trials occurring after successful and failed stops classified as ‘Post Success’ and ‘Post Fail’, respectively. RT in the first Go trial following a stop trial was analysed using a three-way GLMM with fixed factors of Stop Outcome, Condition, and Age Group; participant-level random intercepts, and random slopes by Condition were used.

### Action Cancellation Speed

For each participant, SSRT was estimated for the Finger Tap, Foot Tap, and Step conditions using the integration method ^1^. Per best practice guidelines (Verbruggen *et al.*, 2019), Go omissions (i.e., failure to respond before 1, 1.5 and 2 seconds on Go trials for Finger Tap, Foot Tap and Step conditions, respectively) were replaced with the participant’s slowest RT for that condition. If assumptions of the Horse Race Model were violated; i.e., mean failed stop RT > mean Go trial RT, or when the probability of responding to a stop trial was <25% or >75% for a particular participant/condition the SSRT estimation was excluded from analyses. 5.4% of SSRT values (i.e., 9 out of 168 conditions/participants) were excluded. As SSRT values were normally distributed, a linear mixed model was fitted: *𝑆𝑆𝑅𝑇 = 𝐴𝑔𝑒 𝐺𝑟𝑜𝑢𝑝 x 𝐶𝑜𝑛𝑑𝑖𝑡𝑖𝑜𝑛 + 𝐺𝑒𝑛𝑑𝑒𝑟 + 𝐿𝑎𝑛𝑔𝑢𝑎𝑔𝑒 + Participant ID*

Note that gender was included as a covariate, as previous research has suggested women have faster SSRTs than men ^53^. Significant main and interaction effects were followed up using Bonferroni-adjusted pairwise tests. Additionally, we explored associations between SSRT estimates in the Finger, Foot, and Step conditions using Pearson’s R correlations.

### Modifications to Step Preparation from Action Cancellation Commands

Vertical GRF were calculated from each force plate. In trials with a step, the difference between the GRF of the leading stepping foot and the trailing foot were calculated (ΔGRF) to show the profile of the weight transfer prior to foot-lift. In stop trials without a step (i.e., successful stop trials), the difference between the cued foot (arrow direction) and un-cued foot was calculated as ΔGRF. APAs were identified when there was at least a 5% lateral weight shift that was initiated at least 100ms after the ‘Go’ cue. For each trial, latencies of the APA onset, and peak were determined along with the normalised ΔGRF magnitude for each trial (ΔGRF trial peak amplitude)/(mean ΔGRF peak amplitude of correct steps in CRT condition).

The precise time between the APA peak and the ‘stop’ cue was calculated for each stop trial. With this value plotted on the abscissa and the binary outcome – step inhibition success (0) or failure (1) – on the ordinate axis, a sigmoidal curve was fitted to the data (see Figure 5B). The value at the point of inflection (where the probability of inhibition success is 0.5) was identified as the *Step Threshold* for each participant. This threshold indicates how far into the step preparation process one can be while still being able to use visual feedback to inhibit foot-lift. Analysis of covariance was used to determine the effect of age on step inhibition threshold (relative to APA onset), controlling for differences in mean movement time.

ΔGRF waveforms were examined using Statistical Parametric Mapping (SPM) analysis. SPM allows the use of statistical tests (e.g. ANOVA) to assess for differences in magnitude at each point of a time series ^54^. We used a 2-way ANOVA with the factors of Age Group (Young, Older) and Trial Type (Successful Go, Successful Stop, and Failed Stop), with repeated measures on Trial Type, to test for differences in the force profiles. We also fit a two-way ANOVA comparing Go trials of the CRT and SST blocks (model not shown). A separate model was fit for each timing synchronisation reference: either the ‘Go’ cue, ‘stop’ cue, or peak APA. Bonferroni adjustments were used on post-hoc tests to correct for multiple comparisons.

## Results

### Effects of Age Group, Condition, and Response Type on Response Execution Speed

We firstly compared response times (RTs) between the different groups in the three task conditions in the baseline CRT blocks (where stopping was not a possible response) and SST blocks (where stop signals may occur). There was a significant main effect of Age, *F* (1, Inf) = 52.20, *p* <.001. RT was faster in young adults (*M* = 478ms, 95% CI [458ms, 499ms]) than older adults (*M* = 594ms, 95% CI [570ms, 619ms]). The main effect of Condition was significant, *F* (2, Inf) = 661.70, *p* <.001. RTs were fastest in the Finger Tap condition (*M* = 425ms, 95% CI [410ms, 441ms]), followed by the Foot Tap condition (*M* = 491ms, 95% CI [476ms, 506ms]) and Step condition (*M* = 724ms, 95% CI [701ms, 747ms]). The main effect of Block Type (CRT vs. SST) was significant, *F* (1, Inf) = 58.90, *p* <.001. RTs to ‘go’ signals were faster in the CRT block (*M* = 496ms, 95% CI [484ms, 509ms]) than the SST block (*M* = 572ms, 95% CI [548ms, 596ms]). These main effects are best explained in the context of the significant three-way interaction between Age, Block Type, and Condition, *F* (2, Inf) = 6.04, *p* = .00239 (see Figure 3).

**Figure 3.**
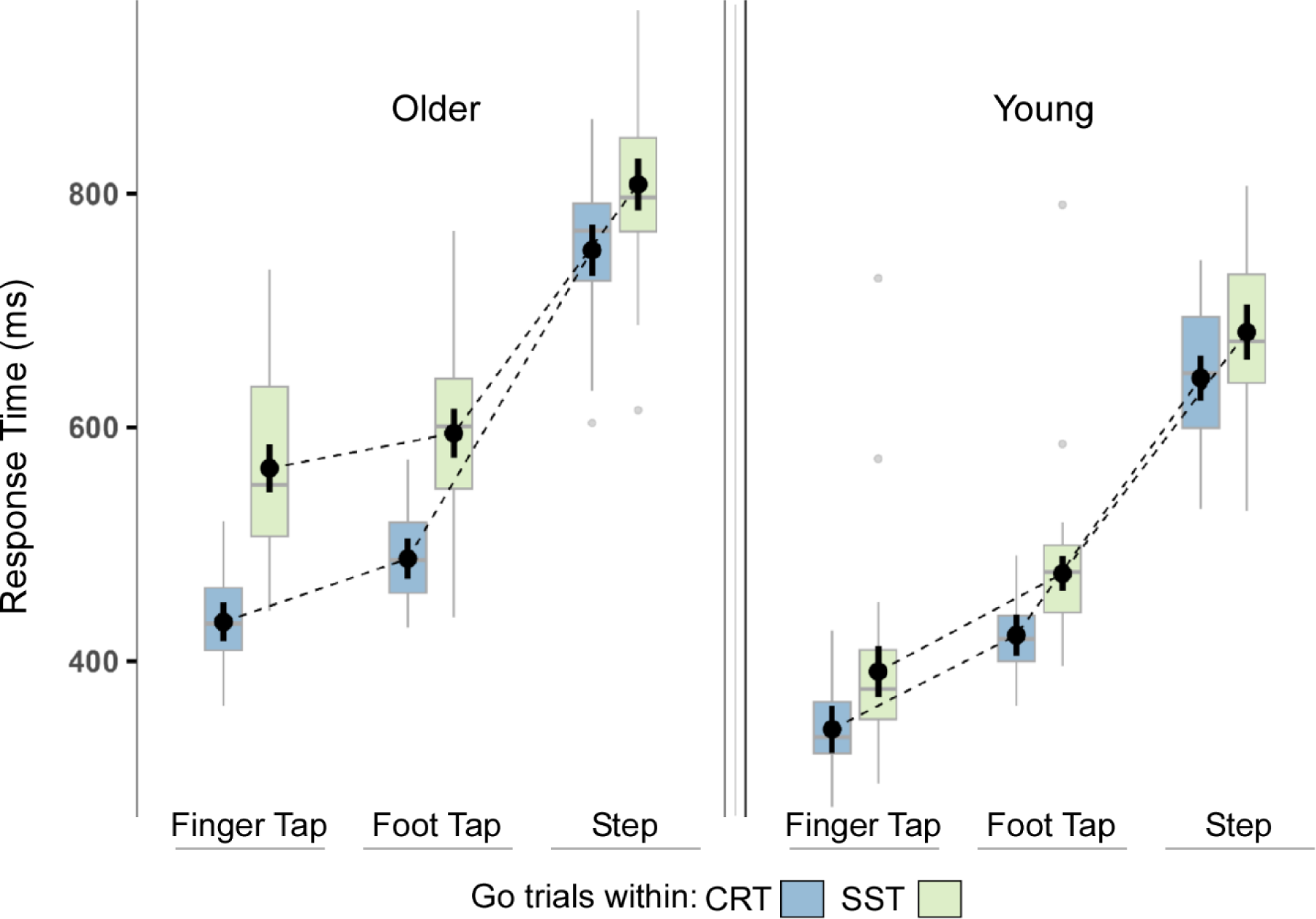
Response Times across conditions. Boxplots represent the median (horizontal line), mean and 95% CI (black circles and black lines), interquartile range (box), maximum (top whisker), and minimum (bottom whisker) of Go RT, stratified by Age Group, Block Type (Blue = CRT, Green = SST) and Condition (Finger Tap, Foot Tap, Step). These plots show a consistent pattern in response speed, with Finger Tap < Foot Tap < Step; CRT < SST; and Young < Older.

Response speed slowing in ‘go’ trials of the SST block (relative to the CRT block) was identified in both age groups, though the magnitude of this was greatest in older adults in the Finger Tap condition. For ‘go’ trials in CRT compared to ‘go’ trials in SST for the Finger Tap, older adults showed a 30% increase in reaction time (95% CI [22%, 39%], ΔRT = 132ms, *z* =7.79, *p* <.001); as opposed to a 14% increase in young adults (95% CI [7%, 23%], ΔRT = 50ms, *z* = 3.83, *p* = <.001). For ‘go’ trials in CRT compared to ‘go’ trials in SST for Foot Tap, older adults showed a 22% increase in RT to ‘go’ signals (95% CI [16%, 29%], ΔRT = 108ms, *z* = 7.11, *p* <.001), as opposed to a 13% increase in young adults (95% CI [6%, 19%], ΔRT = 53ms, *z* = 4.01, *p* <.001). For ‘go’ trials in CRT compared to ‘go’ trials in SST for Step, older adults showed a 7% increase in RT (95% CI [2%, 12%], ΔRT = 55ms, *z* =2.95, *p* = .00315), and young adults showed a 6% increase in RT (95% CI [1%, 12%], ΔRT = 40ms, *z* =2.44, *p* = .0145).

### Effect of Stop Signals on Response Speed in subsequent Go trials

Sequential trial-level slowing in Finger Tap, Foot Tap, and Step conditions following a Stop Signal (SS) trial is depicted in Figure 4 and in Table 3. There was a significant main effect of Trial number post SS across Age Group and Condition, *F* (6, Inf) = 181.75, *p* <.001. Marginal means reveal that RTs were longest immediately after a stop trial (*M* _Post 1_ = 595ms, *SE* = 12.7ms, 95% CI [571ms, 621ms]), and gradually decreased up to 7 trials post SS. Of all levels of the Trial Post SS variable, the Post 7+ level was least affected by slowing. As such, this level was used as the comparator for paired comparison tests.

**Figure 4.**
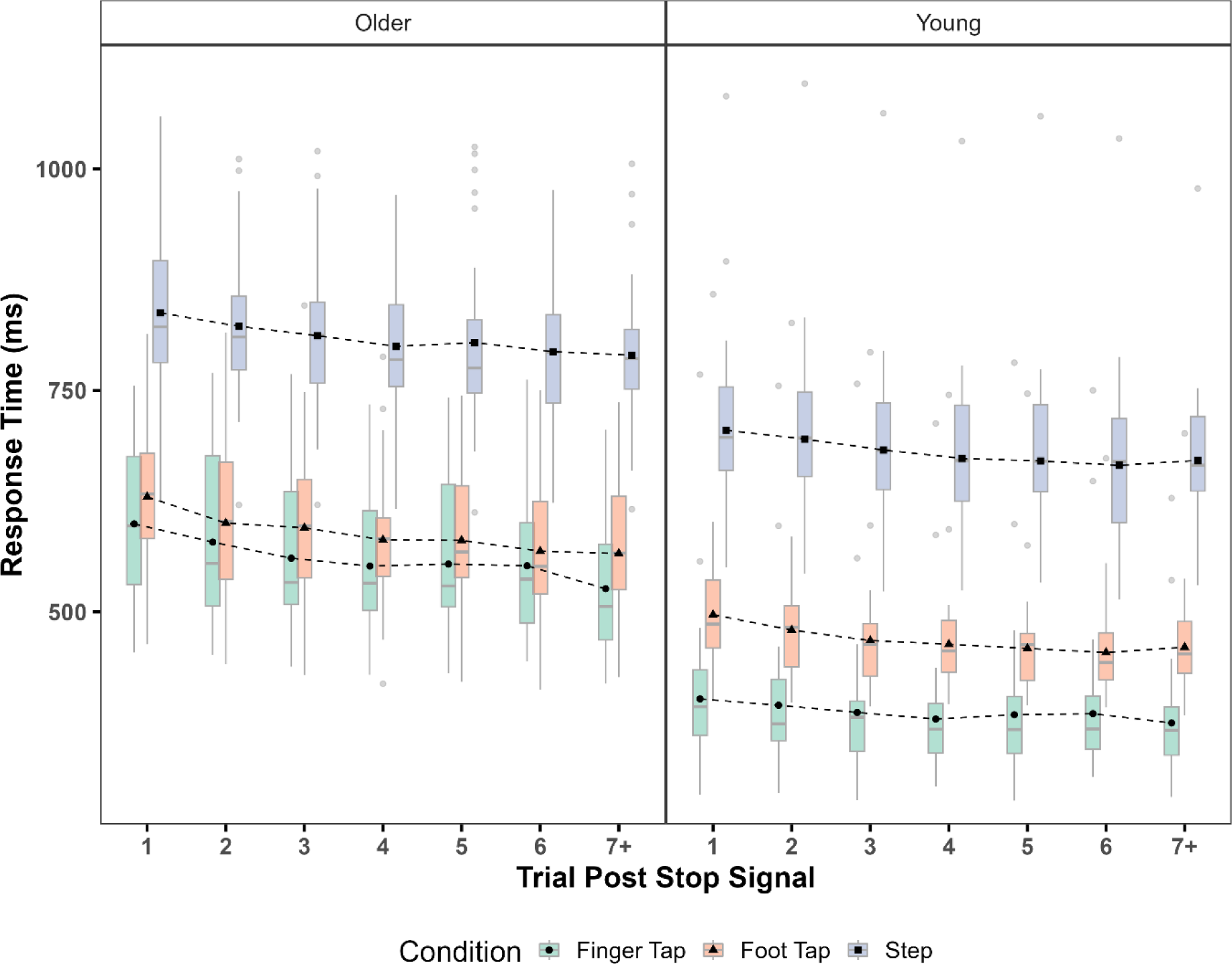
Trial-by-trial changes in response slowing after stop signals. Effect of Age, Condition, and Trial Post Stop Signal on response time. Boxplots represent the median and IQR of RT stratified by Condition (Green = Finger Tap, Orange = Foot Tap, Purple = Step), with group means represented by circles, triangles, and squares respectively. Outliers are represented by grey points immediately above and below the boxplots’ whiskers. This figure indicates RTs were slower for the first trial after a stop signal, and gradually quicken in subsequent Go trials.

**Table 3.**
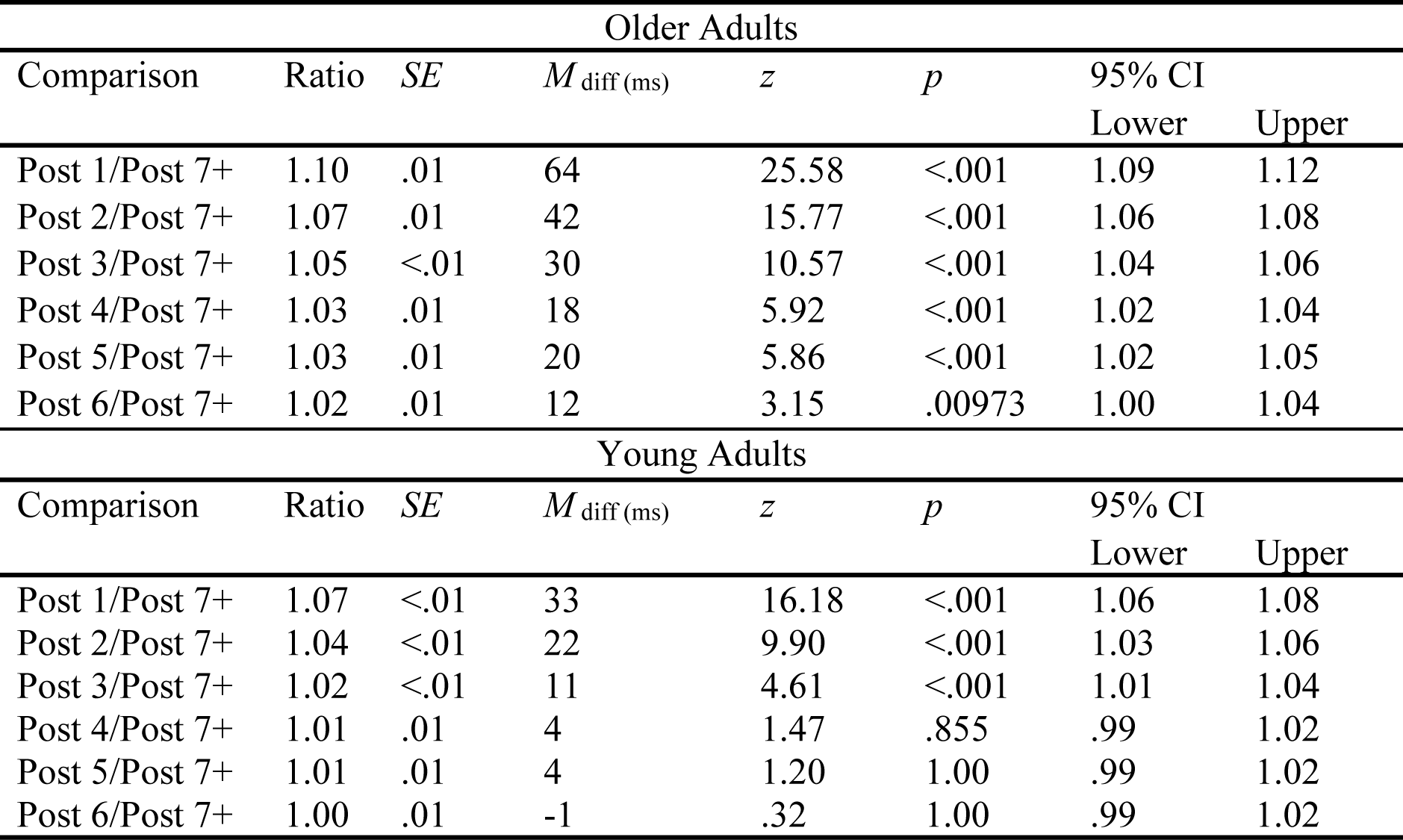
Age Group and RT ratio of trials post stop signal.

| Older Adults |  |  |  |  |  |  |  |
| --- | --- | --- | --- | --- | --- | --- | --- |
| Comparison | Ratio | *SE* | *M*<sub>diff (ms)</sub> | *z* | *p* | 95% CI |  |
|  |  |  |  |  |  | Lower | Upper |
| Post 1/Post 7+ | 1.10 | .01 | 64 | 25.58 | <.001 | 1.09 | 1.12 |
| Post 2/Post 7+ | 1.07 | .01 | 42 | 15.77 | <.001 | 1.06 | 1.08 |
| Post 3/Post 7+ | 1.05 | <.01 | 30 | 10.57 | <.001 | 1.04 | 1.06 |
| Post 4/Post 7+ | 1.03 | .01 | 18 | 5.92 | <.001 | 1.02 | 1.04 |
| Post 5/Post 7+ | 1.03 | .01 | 20 | 5.86 | <.001 | 1.02 | 1.05 |
| Post 6/Post 7+ | 1.02 | .01 | 12 | 3.15 | .00973 | 1.00 | 1.04 |
| Young Adults |  |  |  |  |  |  |  |
| Comparison | Ratio | *SE* | *M*<sub>diff (ms)</sub> | *z* | *p* | 95% CI |  |
|  |  |  |  |  |  | Lower | Upper |
| Post 1/Post 7+ | 1.07 | <.01 | 33 | 16.18 | <.001 | 1.06 | 1.08 |
| Post 2/Post 7+ | 1.04 | <.01 | 22 | 9.90 | <.001 | 1.03 | 1.06 |
| Post 3/Post 7+ | 1.02 | <.01 | 11 | 4.61 | <.001 | 1.01 | 1.04 |
| Post 4/Post 7+ | 1.01 | .01 | 4 | 1.47 | .855 | .99 | 1.02 |
| Post 5/Post 7+ | 1.01 | .01 | 4 | 1.20 | 1.00 | .99 | 1.02 |
| Post 6/Post 7+ | 1.00 | .01 | -1 | .32 | 1.00 | .99 | 1.02 |

The three-way interaction effect between Age Group, Condition, and Trial Post SS, was not significant, *F* (12, Inf) = .1.70, *p =* .0594. However, the two-way interaction between Age Group and Trial Post SS was significant, *F* (6, Inf) = 6.21, *p* <.001 (see Table 3). In older adults, RT was longest on the first trial after a stop (+10% relative to the Post 7+ trial) and decreased with each subsequent trial but remained significant. In contrast, RTs in young adults were elevated for three trials (post 1, post 2, and post 3, relative to post 7+) after a stop signal, but there was no significant difference in RT between the Post 7+ level of the Trial Post SS variable and the Post 4, Post 5, and Post 6 levels.

The two-way interaction between Condition and Trial Post SS was significant, *F* (12, Inf) = 6.40, *p* <.001 (see Table 4). In the Finger Tap condition, the RT for the Post 7+ level of the Trial Post SS variable was significantly longer than all other levels. In the Foot Tap condition, RT slowing persisted for up to four trials after a stop. After this point, RTs were not significantly different to the Post 7+ level. In the Step condition, RT slowing persisted for up to three trials after a stop. After this point, no significant differences were evident relative to the 7+ level.

**Table 4.**
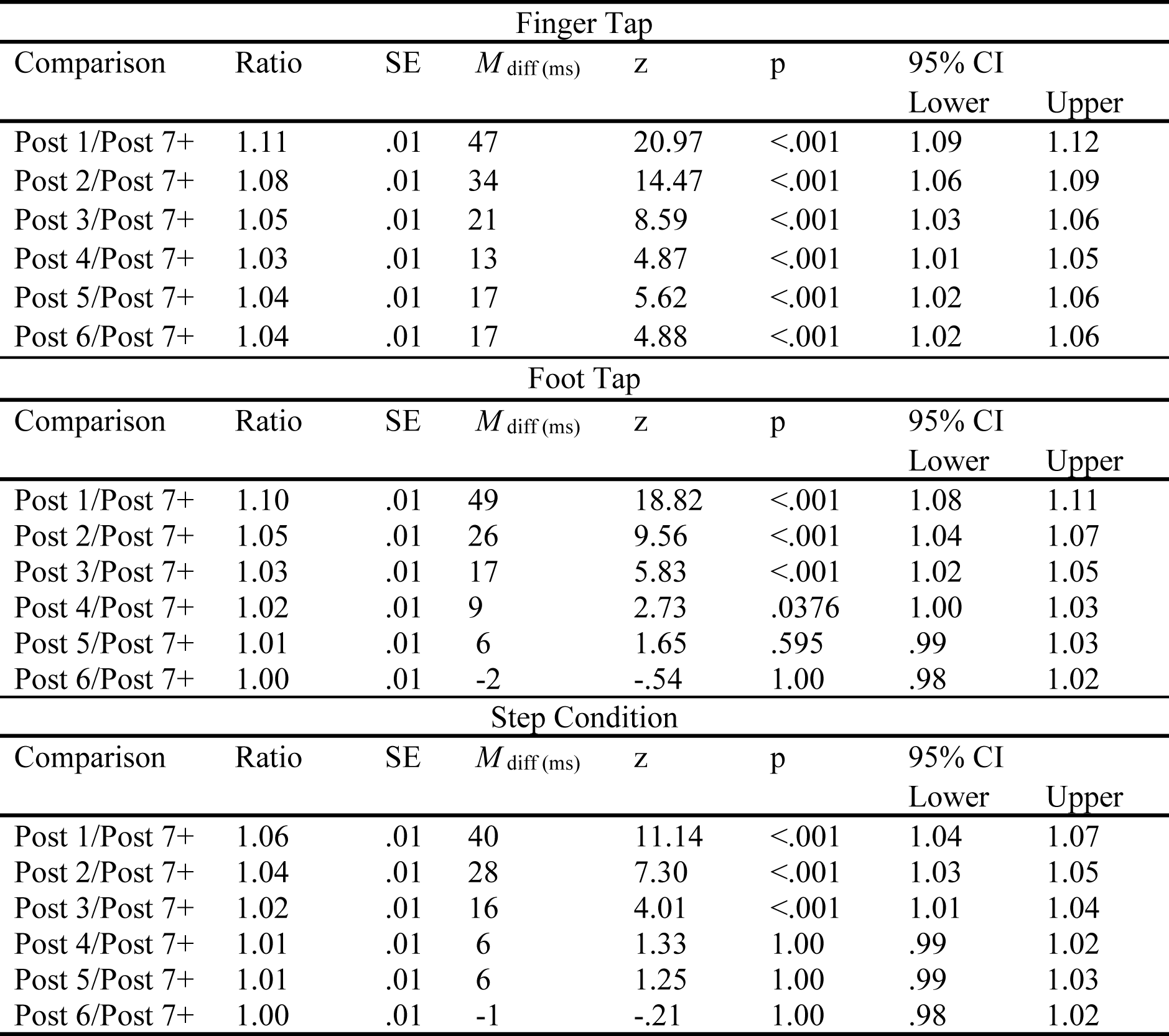
Response modality and RT ratio in trials post stop signal.

| Finger Tap |  |  |  |  |  |  |  |
| --- | --- | --- | --- | --- | --- | --- | --- |
| Comparison | Ratio | SE | $M_{\text{diff (ms)}}$ | z | p | 95% CI | |
|  |  |  |  |  |  | Lower | Upper |
| Post 1/Post 7+ | 1.11 | .01 | 47 | 20.97 | <.001 | 1.09 | 1.12 |
| Post 2/Post 7+ | 1.08 | .01 | 34 | 14.47 | <.001 | 1.06 | 1.09 |
| Post 3/Post 7+ | 1.05 | .01 | 21 | 8.59 | <.001 | 1.03 | 1.06 |
| Post 4/Post 7+ | 1.03 | .01 | 13 | 4.87 | <.001 | 1.01 | 1.05 |
| Post 5/Post 7+ | 1.04 | .01 | 17 | 5.62 | <.001 | 1.02 | 1.06 |
| Post 6/Post 7+ | 1.04 | .01 | 17 | 4.88 | <.001 | 1.02 | 1.06 |
| Foot Tap |  |  |  |  |  |  |  |
| Comparison | Ratio | SE | $M_{\text{diff (ms)}}$ | z | p | 95% CI | |
|  |  |  |  |  |  | Lower | Upper |
| Post 1/Post 7+ | 1.10 | .01 | 49 | 18.82 | <.001 | 1.08 | 1.11 |
| Post 2/Post 7+ | 1.05 | .01 | 26 | 9.56 | <.001 | 1.04 | 1.07 |
| Post 3/Post 7+ | 1.03 | .01 | 17 | 5.83 | <.001 | 1.02 | 1.05 |
| Post 4/Post 7+ | 1.02 | .01 | 9 | 2.73 | .0376 | 1.00 | 1.03 |
| Post 5/Post 7+ | 1.01 | .01 | 6 | 1.65 | .595 | .99 | 1.03 |
| Post 6/Post 7+ | 1.00 | .01 | -2 | -.54 | 1.00 | .98 | 1.02 |
| Step Condition |  |  |  |  |  |  |  |
| Comparison | Ratio | SE | $M_{\text{diff (ms)}}$ | z | p | 95% CI | |
|  |  |  |  |  |  | Lower | Upper |
| Post 1/Post 7+ | 1.06 | .01 | 40 | 11.14 | <.001 | 1.04 | 1.07 |
| Post 2/Post 7+ | 1.04 | .01 | 28 | 7.30 | <.001 | 1.03 | 1.05 |
| Post 3/Post 7+ | 1.02 | .01 | 16 | 4.01 | <.001 | 1.01 | 1.04 |
| Post 4/Post 7+ | 1.01 | .01 | 6 | 1.33 | 1.00 | .99 | 1.02 |
| Post 5/Post 7+ | 1.01 | .01 | 6 | 1.25 | 1.00 | .99 | 1.03 |
| Post 6/Post 7+ | 1.00 | .01 | -1 | -.21 | 1.00 | .98 | 1.02 |

### Effect of Stop Trial Outcome on Subsequent Go Response Speed

We sought to determine whether slowing in the trial immediate after a stop trial was affected by successfully stopping or failing to stop. The three-way interaction between Age Group, Condition, and Stop Outcome was not significant, *F* (2, Inf) = 1.86, *p* = .156. Additionally, neither the two-way interaction between Condition and Stop Outcome nor the main effect of Stop Outcome were significant, *F* (2, Inf) = 2.89, *p* = .0554 and *F* (1, Inf) = <.01, *p* = .955. However, there was a significant two-way interaction between Stop Outcome and Age Group, *F* (2, Inf) = 12.36, *p* <.001. In older adults, RTs were longer after failed than successful stop trials (RT_Post Fail_ - RT_Post Success_ = 9ms, ratio = .99 [95% CI .98, 1.00], *SE* = .01, *z* = −2.50, *p* = .0124); whereas for young adults RTs were longer following successful versus failed stopping (RT_Post Fail_ - RT_Post Success_ = −7ms [95% CI 1.00, 1.03], ratio = 1.01, *SE* = .01, *z* = 2.74, *p* = .0134.

#### Action cancellation speed

Figure 5A shows stopping speed for the different Age Groups and Conditions. SSRT did not differ significantly as a function of Age Group (*F* (1, 63.62) = 1.52, *p* = .222), bilingualism (*F* (4, 60.54) = .69, *p* = .604) or gender (*F* (1, 63.31) = .17, *p* = .684). The two-way interaction between Age Group and Condition was also not significant, *F* (2, 107.99) = .08, *p* = .927. However, there was a significant main effect of Condition (*F* (2, 107.99) = 347.08, *p* <.001). SSRT for the Finger Tap condition was faster than both the Foot Tap condition (*M* _Foot Tap –_ *M* _Finger Tap_ = 102ms, *SE* = 6.06, *t* = 16.79, *p* <.001) and the Step condition (*M* _Step –_ *M* _Finger Tap_ = 156ms, *SE* = 6.02, *t* = 25.93 *p* <.001), and SSRT for the Foot Tap condition was faster than the Step condition (*M* _Step –_ *M* _Foot Tap_ = 54ms, *SE* = 6.09, *t* = 8.91, *p* <.001). SSRT estimates were positively correlated between all conditions: Finger and Foot Tap (*R* = .31, *p* = .033); Finger and Step (*R* = .41, *p* <.001); and Foot Tap and Step (*R* = .58, *p* = <.001).

**Figure 5.**
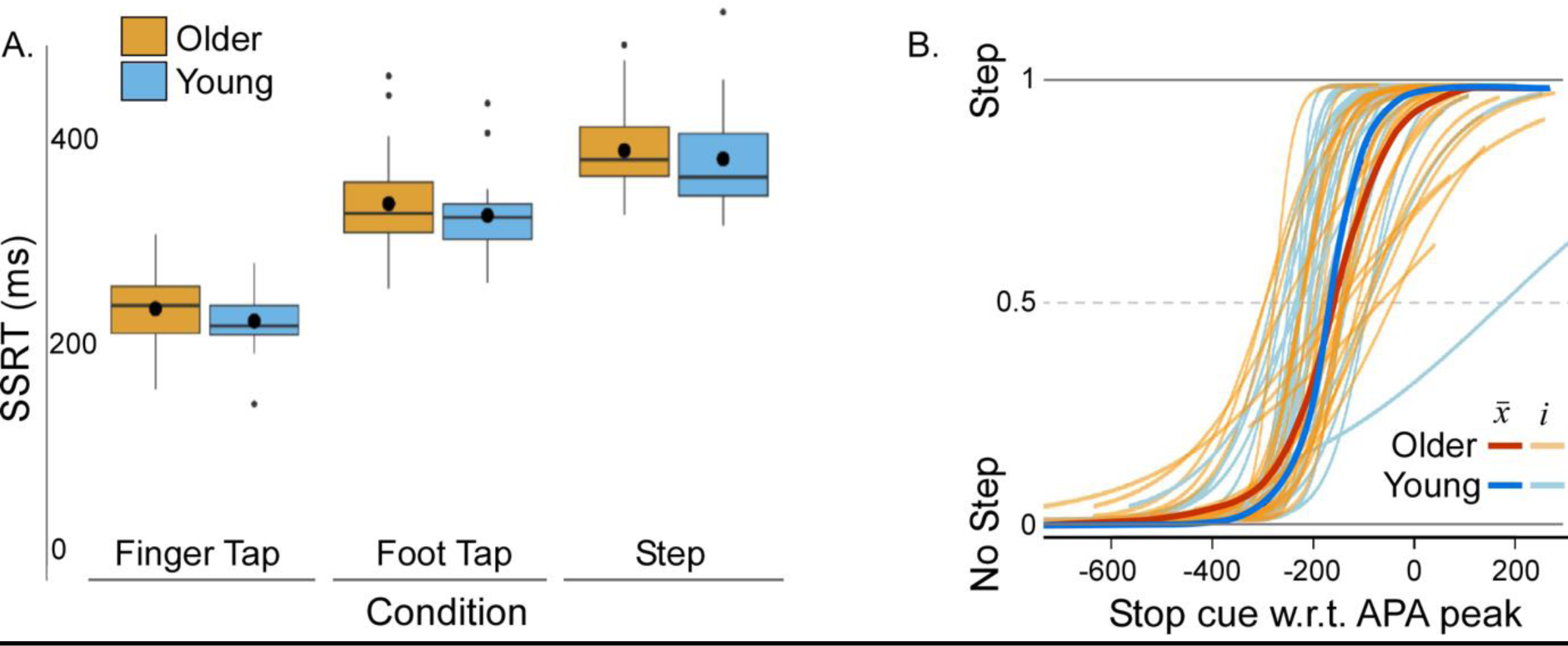
Action cancellation. Boxplots **(A)** represent the median (horizontal line), mean (black circles), interquartile range (box), maximum (top whisker), and minimum (bottom whisker) for SSRT (a single estimate per participant for each of the three conditions). **B.** Step Inhibition Threshold by Age Group depicted as overlaid sigmoid curves. Individual data (*i*) are represented as blue lines (young adults) and yellow lines (older adults). Group mean data are represented as darker blue lines (young adults) and dark red lines (older adults). The x axis represents the timing of the APA peak with respect to the stop signal, and the y axis represents the trial outcome and inhibition threshold (1 = failure to inhibit a step; 0 = successfully inhibited step; 0.5 = step inhibition threshold).

#### Cancellation cues during step preparation

For the Step Condition, the relative weight shifts between the feet (*Δ*GRF waveforms) reflect the underlying mechanisms of step preparation prior to foot-lift. APAs (pre-foot-lift, weight shifts initially toward the side of foot-lift) were apparent on 96.9% of all trials; in young adults, 95.1% of all ‘Go’ trials and 95.25% of ‘Stop’ trials; in older adults, 98.72% of all ‘Go’ trials and 97.76% of all ‘Stop’ trials. The onset time of visual cancellation cues relative to these waveforms and subsequent success or failure in cancelling the foot lift were compared between Age Group within the Step Condition. Step inhibition threshold (Figure 5B) was not significantly different between young and older adults, *F* (1, 52) = 2.37, *p* = .129, indicating that the stop cue could be presented at a similar stage of APA execution and both young and older adults had no difference in the likelihood of cancelling the subsequent step. The *Δ*GRF waveforms for all step trials for all participants are depicted as stacked heatmaps in Figure 6. There is a characteristic weight shift *towards* the stepping foot (APA shown in red) prior to the foot-lift (RT of foot-lift shown as a green dot).

**Figure 6.**
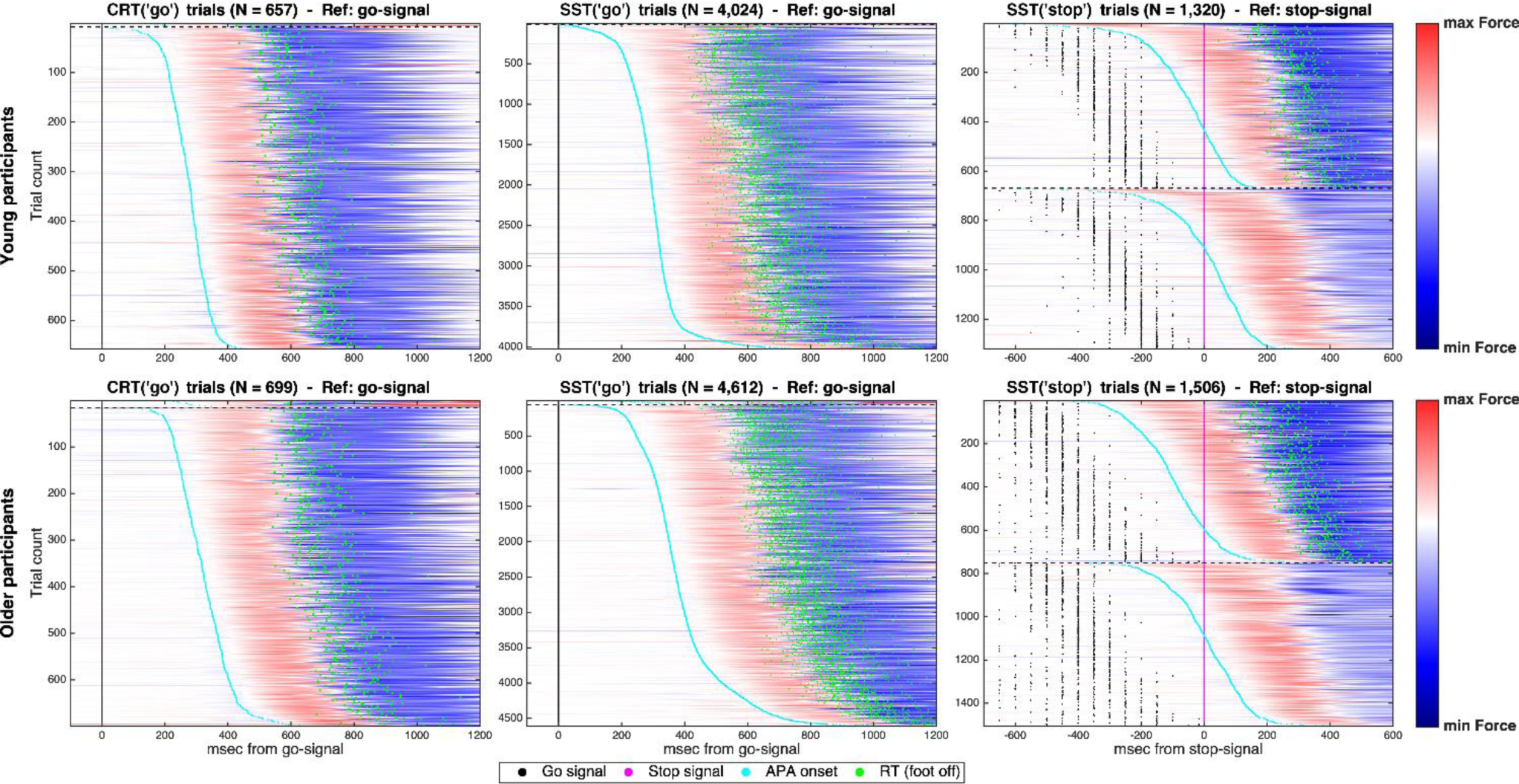
*Δ*GRF for young and older adults. Horizontal lines depict ΔGRF waveforms (leading foot – following foot) as heatmaps for all participants and trials. ΔGRF on ‘Go’ trials of the CRT block (left column), Go trials from the SST block (middle column) and stop trials (right column). Successful stop trials in the right column are shown *below* the horizontal black dotted lines, with failed step cancellation trials (∼50%) represented above this line. The x axis represents time in milliseconds, and the y axis represents trial count (1 coloured horizontal heatmap line = 1 trial). Events within each trial are shown as colour-coded dots according to the legend (bottom of plot). During normal quiet standing (ref: prior to and immediately after the Go signal presentation), there was little to no difference in GRF under each foot (represented by the white colour). During the APA, the vertical ground reactive force initially increases under the leading foot (represented by the colour change to red). When the leading foot is subsequently lifted (green dot), the force drops toward 0 for that leg as the following foot takes the body weight (represented by the colour change in blue). After the second foot-lift, the force difference between the two plates is equalised as no weight is detected when the participant has shifted themselves forward (represented by the colour change back to white toward the end of the many stepping trials).

The average ΔGRF waveforms for Age Group and Trial Type are presented in Figure 7 along with the results of the SPM analyses. APAs were initiated earlier in young than older adults for all trial types (Figure 7, left panel, significant time windows between Young and Older adults), corresponding to faster foot-lift RTs for the young adults.

**Figure 7.**
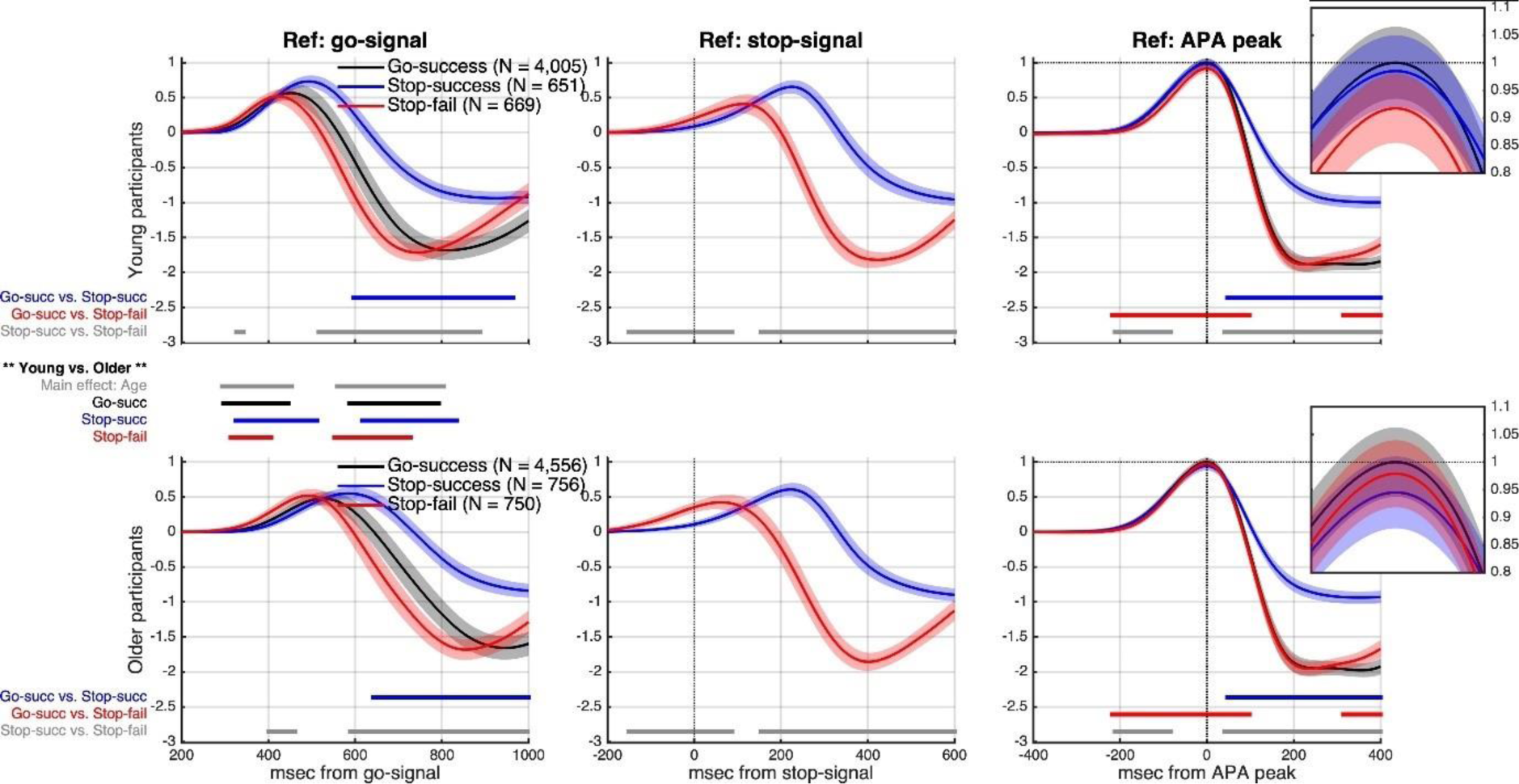
SPM Waveform analysis. Relative changes in vertical ground reaction forces for each trial type, referenced to the go signal (left panel), the stop signal (middle panel), and the peak APA (right panel). Time (ms) is shown on the x axis of each figure. Continuous black (go success within the SST block), blue (stop success), and red (stop fail) waveforms represent changes in mean ΔGRF (normalised to the average of APA peaks across all successful go trials in the SST block), with 95% CIs shown as paler colour bands. ΔGRF ∼0 indicates equal weight between the feet. The first upward inflection of ΔGRF represents the initial weight transfer toward the stepping foot, and the turning point represents the peak APA magnitude, followed by unloading of the stepping foot toward the trailing foot and subsequent foot-lift for the ‘Go-success’ and ‘stop-fail’ trials). The horizontal-coloured lines represent the time points at which the SPM analyses detected statistically significant differences in ΔGRF waveforms from the full factorial 2-way ANOVA (age group, trial type). The lines overlayed inside each plot correspond to significant within-group differences based on post-hoc tests of the interaction between age group and trial type. The horizontal lines presented between the young (top row) and older (bottom row) groups indicate significant between-group differences overall (main effect of age) or in each condition (interaction between age group or trial type).

When the ΔGRF waveforms were synchronised to the presentation of the stop signal (Figure 7, middle panel), the interaction between Age Group and Trial Type was not significant, however APAs were initiated earlier for trials in which participants failed to cancel the step after a stop signal compared to those in which they were successful in cancelling the step.

When *Δ*GRF waveforms were synchronised to APA peak (Figure 7, right panel) there was a significant main effect of Trial Type, as APAs on failed stop trials were faster (reaching APA peak sooner) compared to both go step trials and successful stop trials. The magnitude of the APA peak was smaller for failed stop trials compared to go trials. However there was no significant difference between the size and shape of the APA associated with a normal step response to a go signal, and a step in which an APA was initiated but the foot-lift was successfully aborted (note differences between the ΔGRF waveforms in ‘Go-success’ vs. ‘Stop-success’ trials only occur ∼50ms after the APA peak, but ∼100ms before foot-lift for both young and older groups). There were no significant interactions between Age Group and Trial Type.

## Discussion

This study adapted a stop signal paradigm to evaluate the speed at which older and young adults could initiate and cancel both upper- and lower-limb movements. By using bilateral force plates underfoot, the study applied the stop signal task to voluntary stepping, providing insights into how visually cued action cancellation commands impact postural preparation and step behaviour. This approach has strong ecological relevance, as it may help to better understand age-related impairments in stepping decisions in daily life.

While young adults executed all upper- and lower-limb movements faster than older adults, consistent with prior research showing a general age-related slowing of response speed, a key finding was that the speed to cancel movements was not significantly different between the two age groups. This result supports the notion of independence between motor execution and motor inhibition processes, with the older adults exhibiting less decline in stopping speed compared to execution speed. Age deficits of SSRT identified in other studies tend to be for more complex experimental paradigms which require increased cognitive load (Andrés *et al.*, 2008; Sebastian *et al.*, 2013; Bloemendaal *et al.*, 2016), whereas our results align with prior research in the upper body using a similar, 2-choice response task ^7^.

### Proactive slowing of responses in Older Adults

It is also possible that the similar stopping speeds observed in young and older adults is in part due to proactive slowing when there is an expectation that stopping may be required. This is supported by research showing that when the anticipation of stopping is increased, stopping speed also increases ^55,56^. The current study demonstrates that participants strategically slowed the go response on a trial-wise basis when stops were anticipated, and that the magnitude of slowing was greater for older than young adults. Block-wise analyses of proactive inhibition showed that RT increases were disproportionately greater for older adults in the Finger Tap task (132ms on average, as opposed to 50ms in young adults). This suggests that older adults’ stopping ability was supported by their ability to make proactive adjustments to the speed of the go process. Neuroimaging research indicates that proactive control primes the stopping network. An fMRI-based SST study indicates that frontal-parietal regions thought to be involved in stopping have increased activation on trials with potential to elicit proactive inhibition (“uncertain go” trials, analogous to Go trials of the SST block in the current experiment) relative to “certain Go” trials that do not elicit stopping expectation (analogous to trials of the CRT condition) ^55^. Anticipation of stopping also engages the ‘indirect’ corticobasal pathway, regions typically involved in response inhibition ^11^.

Consistent with previous SST research, we demonstrate that the magnitude of slowing after stop signals decreases with subsequent trials after a stop (Emeric et al., 2007). However, the post-stop slowing of go responses was maintained in older adults for up to six trials after a stop, as opposed to three trials after a stop in young adults. This likely reflects a strategic process enacted when the participants priority switches from fast responding to preparing for a possible stop, known as the goal-priority hypothesis ^14^.

The post-stop slowing of responses occurred after both successful and failed action cancellation, consistent with prior research (Reiger & Geigel, 1999; Bissett & Logan, 2011). We show that the magnitude of slowing in each case was equivalent. In other words, while trial RT was at least partially dependent on its serial position; the outcome of the preceding stop trial was irrelevant. Whereas previous studies of post-signal slowing have manipulated participant expectancies of a stop, in the current study the likelihood of sequential stop signals was random, and it was entirely possible for two (or more) stop signals to occur in a row. The uncertainty of stopping likelihood seems to have engendered a more cautious response style in older than young adults, which manifested as slow RTs that persisted long after the appearance of a stop trial.

Slowing after stop signals was most persistent in the Finger Tap condition (enduring up to six trials after a stop), and least persistent in the step condition (enduring up to three trials after a stop). This is directly contrary to the hypotheses, which anticipated that proactive slowing would be most prominent during the step condition. To our knowledge, this is the first study to explore inhibitory after-effects in the lower limb. One explanation was that participants respond cautiously on both CRT and SST Go trials of the Step condition regardless of whether a stop was anticipated, because they were focussed on maintaining stability. With a slow base response time in stepping, additional proactive slowing provides minimal advantage to stopping speed. An alternative explanation points to the *type* of motor control required and its execution speed. As tasks progressed from using fine to gross motor skill, Go trial RT *increased*, and the magnitude and persistence of post-stop slowing *decreased.* Stop signals on the Finger Tap ask may have been particularly salient because movements that were faster and less effortful to initiate were more difficult to stop. When stop signals were presented, participants slowed down strategically to increase the likelihood of stopping on the subsequent trial, and RTs were gradual to recover. Previously, it has been shown that block-wise proactive slowing can be suppressed by introducing automated warnings after slow Go Trial RTs ^57^. Future research should examine the effect of warnings on the persistence of inhibitory after-effects and could elucidate whether they are a conscious strategy.

### Decoupling of the APA from the Step

Before the foot-lift, postural weight shifts toward the stepping foot were observed on almost all trials. Prior studies have defined stepping APAs as the centre of pressure (CoP) shifts that occur before the action of the prime mover: lifting the foot off the ground, thus classifying foot-lift as the onset of voluntary movement ^58,59^. The findings of this study revealed that the magnitude and speed of the APA did not differ between go trials and stop trials in which an APA was initiated but successfully aborted. However, approximately 40ms after the peak loading of the stepping foot, and as the centre of mass was shifting toward the stance leg there was significant divergence between the ground reaction forces of go stepping trials and successfully cancelled stepping trials (Figure 7). This suggests that while the initial phase of the APA (loading of the step foot) is a feedforward, ballistic process, later stages are amenable to feedback from visual stimuli. This permits a decoupling of the foot-lift from the step preparation phase, even when an APA has been triggered. The results showed that healthy older adults (who did not have trouble with falling or concerns about their balance), were able to abort the foot-lift phase of their step when visual stop cues were presented at similar stages of the APA to young adults.

The Horse Race Model of the SST ^60^ provides insight into why APA on certain stop trials were able to be cancelled, whereas others resulted in an incorrect forward step. Specifically, APAs that are initiated earlier in response to the ‘go’ signal, and with greater speed tend to lead to failure of step cancellation, as the stop signal is presented closer to the point of no return.

Moderate correlations were observed between SSRTs for the finger and both lower-limb SSRTs, likely reflecting shared neural circuits involved in global motor inhibition across the body. However, stronger correlations were found between the stopping speed of the foot tap and step initiation, suggesting the involvement of more limb-specific mechanisms for lower-limb motor control. This distinction may reflect the closer functional and anatomical relationship between these lower-limb actions. Prior ageing research has used correlational designs that explore the relationship between upper-limb inhibitory control and lower limb mobility ^61–63^. Given only the moderate correlations found between SSRT at the finger and during stepping revealed in this study, future studies should focus on whether measuring SSRT during a step is more indicative of falls risk, than measuring SSRT in the finger in populations at risk of falling.

### Limitations

This study used many trials (240 per condition) to provide an accurate assessment of action cancellation; however, this robust approach resulted in long testing times (∼2.5 hours total). Whilst accommodations were made to reduce cognitive fatigue, and conditions were counterbalanced to reduce order effects, fatigue may have affected stopping ability and RT. This research would need to be adapted and shortened to explore areas of strong clinical relevance, such as whether stopping speed on the SST declines in older adults with neurodegenerative conditions. Additionally, a fruitful avenue for future research would be to examine proactive and reactive inhibition during stepping in older adults with high risk of falls.

### Conclusions

This study suggests that stopping speed was not affected by healthy ageing but does indicate that ageing affects the extent to which proactive strategies are employed in contexts that *may* require stopping. Additionally, these results demonstrate that whilst initial phases of postural preparation for a step appear to be ballistic, inhibitory motor commands can decouple APA and foot-lift stages of a step with similar success in both young and healthy older adults.

## Acknowledgements

The researchers acknowledge the contribution of Olivia Hannon to the data collection process. We would also like to extend our thanks to the people who kindly volunteered their time to participate in this experiment.

## Author Contributions

RH, RStG, RP, and MH conceived the experiment. RH, RStG, and MW collected data. RH, RStG and SS conducted formal analysis. RStG and MH supervised the project. RH and RStG wrote the initial draft of the manuscript, and all authors reviewed and edited the final manuscript.

## Data Availability

The data that support the findings of this study are available from the corresponding author, R StG upon reasonable request.

## References

1 Verbruggen, F. et al. A consensus guide to capturing the ability to inhibit actions and impulsive behaviors in the stop-signal task. Elife 8, doi:10.7554/eLife.46323 (2019).

2 Deary, I. J. & Der, G. Reaction Time, Age, and Cognitive Ability: Longitudinal Findings from Age 16 to 63 Years in Representative Population Samples. Aging, Neuropsychology, and Cognition 12, 187–215, doi:10.1080/13825580590969235 (2005).

3 Lord, S. R. & Fitzpatrick, R. C. Choice stepping reaction time: a composite measure of falls risk in older people. J Gerontol A Biol Sci Med Sci 56, M627–632, doi:10.1093/gerona/56.10.m627 (2001).

4 Andrés, P., Guerrini, C., Phillips, L. H. & Perfect, T. J. Differential Effects of Aging on Executive and Automatic Inhibition. Developmental Neuropsychology 33, 101–123, doi:10.1080/87565640701884212 (2008).

5 Bloemendaal, M. et al. Contrasting neural effects of aging on proactive and reactive response inhibition. Neurobiology of Aging 46, 96–106, doi:10.1016/j.neurobiolaging.2016.06.007 (2016).

6 Sebastian, A. et al. Differential effects of age on subcomponents of response inhibition. Neurobiology of Aging 34, 2183–2193, doi:10.1016/j.neurobiolaging.2013.03.013 (2013).

7 Williams, B. R., Ponesse, J. S., Schachar, R. J., Logan, G. D. & Tannock, R. Development of inhibitory control across the life span. Dev Psychol 35, 205–213, doi:10.1037//0012-1649.35.1.205 (1999).

8 Smittenaar, P. et al. Proactive and Reactive Response Inhibition across the Lifespan. PLOS ONE 10, e0140383, doi:10.1371/journal.pone.0140383 (2015).

9 Bedard, A.-C. et al. The Development of Selective Inhibitory Control Across the Life Span. Developmental Neuropsychology 21, 93–111, doi:10.1207/s15326942dn2101_5 (2002).

10 Van De Laar, M., Van Den Wildenberg, W., van Boxtel, G. & van der Molen, M. Lifespan Changes in Global and Selective Stopping and Performance Adjustments. Frontiers in Psychology 2, doi:10.3389/fpsyg.2011.00357 (2011).

11 Aron, A. R. From Reactive to Proactive and Selective Control: Developing a Richer Model for Stopping Inappropriate Responses. Biological Psychiatry 69, e55–e68, doi:10.1016/j.biopsych.2010.07.024 (2011).

12 Meyer, H. C. & Bucci, D. J. Neural and behavioral mechanisms of proactive and reactive inhibition. Learning & Memory 23, 504–514, doi:10.1101/lm.040501.115 (2016).

13 Verbruggen, F., Logan, G. D., Liefooghe, B. & Vandierendonck, A. Short-term aftereffects of response inhibition: repetition priming or between-trial control adjustments? J Exp Psychol Hum Percept Perform 34, 413–426, doi:10.1037/0096-1523.34.2.413 (2008).

14 Bissett, P. G. & Logan, G. D. Post-stop-signal slowing: Strategies dominate reflexes and implicit learning. Journal of Experimental Psychology: Human Perception and Performance 38, 746–757, doi:10.1037/a0025429 (2012).

15 Li, C.-s. R., et al. Neural Correlates of Post-error Slowing during a Stop Signal Task: A Functional Magnetic Resonance Imaging Study. Journal of Cognitive Neuroscience 20, 1021–1029, doi:10.1162/jocn.2008.20071 (2008).

16 Verbruggen, F. & Logan, G. D. Long-term aftereffects of response inhibition: Memory retrieval, task goals, and cognitive control. Journal of Experimental Psychology: Human Perception and Performance 34, 1229–1235, doi:10.1037/0096-1523.34.5.1229 (2008).

17 Yu, C.-C., Muggleton, N. G., Chen, C.-Y., Ko, C.-H. & Liu, S. The comparisons of inhibitory control and post-error behaviors between different types of athletes and physically inactive adults. PLOS ONE 16, e0256272, doi:10.1371/journal.pone.0256272 (2021).

18 Mirelman, A. et al. Executive Function and Falls in Older Adults: New Findings from a Five-Year Prospective Study Link Fall Risk to Cognition. PLoS ONE 7, e40297, doi:10.1371/journal.pone.0040297 (2012).

19 Anstey, K. J., Wood, J., Kerr, G., Caldwell, H. & Lord, S. R. Different cognitive profiles for single compared with recurrent fallers without dementia. Neuropsychology 23, 500 (2009).

20 Koo, D.-K. & Kwon, J.-W. Biomechanical Analysis of Unplanned Gait Termination According to a Stop-Signal Task Performance: A Preliminary Study. Brain Sciences 13, 304, doi:10.3390/brainsci13020304 (2023).

21 England, D. et al. Relationship between Speed of Response Inhibition and Ability to Suppress a Step in Midlife and Older Adults. Brain Sciences 11, 643, doi:10.3390/brainsci11050643 (2021).

22 Bissett, P. G. et al. Generalized motor inhibitory deficit in Parkinson’s disease patients who freeze. Journal of Neural Transmission 122, 1693–1701, doi:10.1007/s00702-015-1454-9 (2015).

23 Tabu, H., Mima, T., Aso, T., Takahashi, R. & Fukuyama, H. Common inhibitory prefrontal activation during inhibition of hand and foot responses. Neuroimage 59, 3373–3378, doi:10.1016/j.neuroimage.2011.10.092 (2012).

24 Tatz, J. R., Soh, C. & Wessel, J. R. Common and Unique Inhibitory Control Signatures of Action-Stopping and Attentional Capture Suggest That Actions Are Stopped in Two Stages. The Journal of Neuroscience 41, 8826–8838, doi:10.1523/jneurosci.1105-21.2021 (2021).

25 Bolton, D. A. E. & Richardson, J. K. Inhibitory Control and Fall Prevention: Why Stopping Matters. Frontiers in Neurology 13, doi:10.3389/fneur.2022.853787 (2022).

26 Schoene, D., Delbaere, K. & Lord, S. R. Impaired Response Selection During Stepping Predicts Falls in Older People-A Cohort Study. J Am Med Dir Assoc 18, 719–725, doi:10.1016/j.jamda.2017.03.010 (2017).

27 Raud, L., Westerhausen, R., Dooley, N. & Huster, R. J. Differences in unity: The go/no-go and stop signal tasks rely on different mechanisms. NeuroImage 210, 116582, doi:10.1016/j.neuroimage.2020.116582 (2020).

28 Schoene, D., Smith, S. T., Davies, T. A., Delbaere, K. & Lord, S. R. A Stroop Stepping Test (SST) using low-cost computer game technology discriminates between older fallers and non-fallers. Age Ageing 43, 285–289, doi:10.1093/ageing/aft157 (2014).

29 Pelicioni, P. H., Lord, S. R., Okubo, Y., Sturnieks, D. L. & Menant, J. C. People with Parkinson’s disease exhibit reduced cognitive and motor cortical activity when undertaking complex stepping tasks requiring inhibitory control. Neurorehabilitation and Neural Repair 34, 1088–1098 (2020).

30 Sparto, P. J. et al. Postural adjustment errors reveal deficits in inhibition during lateral step initiation in older adults. Journal of Neurophysiology 109, 415–428 (2013).

31 Sparto, P. J. et al. Postural adjustment errors during lateral step initiation in older and younger adults. Exp Brain Res 232, 3977–3989, doi:10.1007/s00221-014-4081-z (2014).

32 Magnard, J. et al. Perceptual Inhibition Is Not a Specific Component of the Sensory Integration Process Necessary for a Rapid Voluntary Step Initiation in Healthy Older Adults. The Journals of Gerontology: Series B 75, 1921–1929, doi:10.1093/geronb/gbz060 (2019).

33 Kwag, E., Bachmann, D., Kim, K., Komnik, I. & Zijlstra, W. Effects of cognitive inhibition preceding voluntary step responses to visual stimuli in young and older adults. The Journals of Gerontology, Series B: Psychological Sciences and Social Sciences 79, gbae006 (2024).

34 Uemura, K., Haruta, M. & Uchiyama, Y. Age differences in reactive strategies and execution time during choice stepping with visual interference. Eur J Appl Physiol 116, 1053–1062, doi:10.1007/s00421-015-3221-x (2016).

35 Coelho, D. B. et al. Frontal Hemodynamic Response During Step Initiation Under Cognitive Conflict in Older and Young Healthy People. The Journals of Gerontology: Series A 76, 216–223 (2021).

36 Kwag, E., Komnik, I., Bachmann, D. & Zijlstra, W. Motor inhibition during voluntary gait initiation in young and older adults. Scientific Reports 14, doi:10.1038/s41598-024-79790-5 (2024).

37 Potocanac, Z., Smulders, E., Pijnappels, M., Verschueren, S. & Duysens, J. Response inhibition and avoidance of virtual obstacles during gait in healthy young and older adults. Hum Mov Sci 39, 27–40, doi:10.1016/j.humov.2014.08.015 (2015).

38 Nissan, M. & Whittle, M. W. Initiation of gait in normal subjects: a preliminary study. Journal of Biomedical Engineering 12, 165–171, 10.1016/0141-5425(90)90139-E (1990).

39 Lyon, I. N. & Day, B. L. Control of frontal plane body motion in human stepping. Experimental Brain Research 115, 345–356, doi:10.1007/pl00005703 (1997).

40 Massion, J. Movement, posture and equilibrium: interaction and coordination. Prog Neurobiol 38, 35–56, doi:10.1016/0301-0082(92)90034-c (1992).

41 Bancroft, M. J. & Day, B. L. The Throw-and-Catch Model of Human Gait: Evidence from Coupling of Pre-Step Postural Activity and Step Location. Front Hum Neurosci 10, 635, doi:10.3389/fnhum.2016.00635 (2016).

42 St George, R. J., Carlson-Kuhta, P., King, L. A., Burchiel, K. J. & Horak, F. B. Compensatory stepping in Parkinson’s disease is still a problem after deep brain stimulation randomized to STN or GPi. Journal of Neurophysiology 114, 1417–1423, doi:10.1152/jn.01052.2014 (2015).

43 King, L. A., St George, R. J., Carlson-Kuhta, P., Nutt, J. G. & Horak, F. B. Preparation for compensatory forward stepping in Parkinson’s disease. Arch Phys Med Rehabil 91, 1332–1338, doi:10.1016/j.apmr.2010.05.013 (2010).

44 Sturnieks, D. L., St George, R. & Lord, S. R. Balance disorders in the elderly. Neurophysiologie Clinique/Clinical Neurophysiology 38, 467–478, 10.1016/j.neucli.2008.09.001 (2008).

45 Yardley, L. et al. Development and initial validation of the Falls Efficacy Scale-International (FES-I). Age and Ageing 34, 614–619, doi:10.1093/ageing/afi196 (2005).

46 Marian, V., Blumenfeld, H. K. & Kaushanskaya, M. The Language Experience and Proficiency Questionnaire (LEAP-Q): Assessing Language Profiles in Bilinguals and Multilinguals. Journal of Speech, Language, and Hearing Research 50, 940–967, doi:doi:10.1044/1092-4388(2007/067) (2007).

47 Bialystok, E., Martin, M. M. & Viswanathan, M. Bilingualism across the lifespan: The rise and fall of inhibitory control. International Journal of Bilingualism 9, 103–119, doi:10.1177/13670069050090010701 (2005).

48 Bialystok, E., Craik, F. I. M. & Ryan, J. Executive control in a modified antisaccade task: Effects of aging and bilingualism. Journal of Experimental Psychology: Learning, Memory, and Cognition 32, 1341–1354, doi:10.1037/0278-7393.32.6.1341 (2006).

49 Verbruggen, F., Logan, G. D. & Stevens, M. A. STOP-IT: Windows executable software for the stop-signal paradigm. Behavior Research Methods 40, 479–483, doi:10.3758/brm.40.2.479 (2008).

50 Lo, S. & Andrews, S. To transform or not to transform: using generalized linear mixed models to analyse reaction time data. Frontiers in Psychology 6, doi:10.3389/fpsyg.2015.01171 (2015).

51 Singmann, H. & Kellen, D. 4–31 (2019).

52 Barr, D. J., Levy, R., Scheepers, C. & Tily, H. J. Random effects structure for confirmatory hypothesis testing: Keep it maximal. Journal of Memory and Language 68, 255–278, 10.1016/j.jml.2012.11.001 (2013).

53 Rubia, K. et al. Effects of age and gender on neural networks of motor response inhibition: From adolescence to mid-adulthood. NeuroImage 83, 690–703, 10.1016/j.neuroimage.2013.06.078 (2013).

54 Pataky, T. C. One-dimensional statistical parametric mapping in Python. Computer Methods in Biomechanics and Biomedical Engineering 15, 295–301, doi:10.1080/10255842.2010.527837 (2012).

55 Chikazoe, J. et al. Preparation to Inhibit a Response Complements Response Inhibition during Performance of a Stop-Signal Task. The Journal of Neuroscience 29, 15870, doi:10.1523/JNEUROSCI.3645-09.2009 (2009).

56 Doekemeijer, R. A., Dewulf, A., Verbruggen, F. & Boehler, C. N. Proactively Adjusting Stopping: Response Inhibition is Faster when Stopping Occurs Frequently. Journal of Cognition, doi:10.5334/joc.264 (2023).

57 Weber, S., Salomoni, S. E., Kilpatrick, C. & Hinder, M. R. Dissociating attentional capture from action cancellation during the inhibition of bimanual movement. Psychophysiology, doi:10.1111/psyp.14372 (2023).

58 Schoene, D., Lord, S. R., Verhoef, P. & Smith, S. T. A Novel Video Game–Based Device for Measuring Stepping Performance and Fall Risk in Older People. Archives of Physical Medicine and Rehabilitation 92, 947–953, 10.1016/j.apmr.2011.01.012 (2011).

59 Lepers, R. & Brenière, Y. The role of anticipatory postural adjustments and gravity in gait initiation. Experimental Brain Research 107, doi:10.1007/bf00228023 (1995).

60 Logan, G. D. & Cowan, W. B. On the ability to inhibit thought and action: A theory of an act of control. Psychological Review 91, 295–327, doi:10.1037/0033-295X.91.3.295 (1984).

61 Caetano, M. J. D. et al. Sensorimotor and Cognitive Predictors of Impaired Gait Adaptability in Older People. J Gerontol A Biol Sci Med Sci 72, 1257–1263, doi:10.1093/gerona/glw171 (2017).

62 Kearney, F. C., Harwood, R. H., Gladman, J. R. F., Lincoln, N. & Masud, T. The Relationship between Executive Function and Falls and Gait Abnormalities in Older Adults: A Systematic Review. Dementia and Geriatric Cognitive Disorders 36, 20–35, doi:10.1159/000350031 (2013).

63 Montero-Odasso, M. & Speechley, M. Falls in Cognitively Impaired Older Adults: Implications for Risk Assessment And Prevention. Journal of the American Geriatrics Society 66, 367–375, doi:10.1111/jgs.15219 (2018).

